# Overwinter roost activity patterns in response to environmental conditions along a rural-urban gradient

**DOI:** 10.64898/2026.10.03.756417

**Authors:** Kelly A. Williams, Jacob E. Morgan, Susan M. Lyons

## Abstract

1. Organisms rely on environmental cues, such as moonlight illumination and weather, to modulate nocturnal behaviors that reduce predation risk and conserve energy during demanding periods. Urbanization can alter the reliability of cues, potentially impairing organisms’ ability to respond appropriately with consequences in increasingly urbanized landscapes.
2. We leveraged the Motus Wildlife Tracking System to identify roost initiation, roost departure and nocturnal restlessness of dark-eyed juncos (*Junco hyemalis*) overwintering along an urban to rural gradient to determine if activity patterns are affected by weather, moonlight illumination, and urbanization.
3. Nocturnal behavior of overwintering juncos varied with environmental factors, including moonlight illumination and temperature. However, the strength of these responses differed among sites, suggesting that natural cues of moonlight illumination were masked in urban areas, where individuals were more responsive to temperature.
4. Our results suggest that urbanization weakens birds’ responsiveness to natural light cues, likely because artificial lighting reduces the visibility of moonlight; representing a key mechanism by which human-altered landscapes can disrupt nocturnal ecology in birds.

## 1. INTRODUCTION

Organisms have evolved physiological and behavioral responses to environmental cues that modulate daily activity cycles (Davidson and Menaker 2003, Ashton et al. 2022, Oosthuizen and Bennett 2022) and support survival and reproductive success (Bradshaw and Holzapfel 2010, Shinomiya et al. 2023, Vieira et al. 2025). A critical component of the daily cycle is nocturnal activity. In diurnal species, foraging and communication are reduced at night, and replaced with nocturnal behaviors that reduce predation risks, increase energy conservation, protect from unfavorable weather conditions, and promote sleep (Webb and Rogers 1988, Tisdale et al. 2018, Ferretti et al. 2019, Neumann et al. 2022). To regulate nocturnal behaviors, organisms use cues related to moonlight illumination, weather, predator density, presence of conspecifics, and roost site availability (Rubolini et al. 2015, Schlicht and Kempenaers 2020, Nomano et al. 2021, Allen 2025, Lowney and Thomson 2026). During winter, shorter days reduce the time available for foraging (Carrascal et al. 2012), while colder temperatures and inclement weather increase thermoregulatory and energetic demands (Webb and Rogers 1988, Stager et al. 2020, Boyer and MacDougall-Shackleton 2020, Li et al. 2022). Given the importance of environmental cues during winter, nocturnal behaviors, including the timing of roost initiation, nocturnal restlessness, and roost departure, may be sensitive to environmental changes, including urbanization.

Urbanization has resulted in multifaceted environmental changes that impact how organisms structure daily cycles. Singing behavior, roost timing, and roost site selection often shift in urban compared to rural environments, driven by co-occurring features of anthropogenic development including artificial light at night (ALAN), noise pollution, urban heat island effects, and changes in predation pressure and food availability (Marín-Gómez and MacGregor-Fors 2021, Chen et al. 2023, Dougherty et al. 2024). ALAN is associated with earlier dawn singing and extended activity into nighttime hours, effectively lengthening active periods at the expense of rest (Kumar et al. 2021, Marín-Gómez 2022, Godet et al. 2026). Anthropogenic noise masks acoustic signals used for communication and predator detection (Zhou et al. 2019, Tilgar et al. 2022). Collectively, these patterns indicate that urbanization directly alters nocturnal behaviors.

Urbanization can alter how responsive organisms are to natural environmental cues that govern nocturnal behaviors. Light pollution can homogenize nighttime light levels and reduce the contribution of lunar cycling on a bird’s activity patterns (Amichai and Kronfeld-Schor 2019, Evens et al. 2023). Similarly, differences in resource availability or buffered thermal conditions can alter activity budgets (Leveau 2018). As environmental variation is masked, unmasked, or homogenized by urban conditions, organisms may become more or less responsive to cues that were previously relied on.

Much of what is known about how environmental conditions shape roost timing and nocturnal activity comes from captive studies, where cues such as light, temperature, and photoperiod can be manipulated independently (Aschoff and Von Goetz 1989, Lehmann et al. 2012, Litt-Jukes and Morrissey 2026). These studies have been valuable in identifying the environmental cues that influence nocturnal behaviors (Kumar et al. 2021, Zuo et al. 2023, Tiwari et al. 2026). However, free-living birds often experience these conditions synergistically. How environmental variation shapes roost initiation, nocturnal activity, and roost departure in free-living birds across urban-to-rural gradients remains less understood.

We used remote sensing technology within the Motus Wildlife Tracking System (Motus) to determine how nocturnal behavior in Dark-eyed Juncos (*Junco hyemalis*) varied in response to environmental variation across 6 sites along an urban, suburban and rural gradient during winter. We measured roost initiation, roost departure, and nocturnal restlessness. We tested the hypotheses that roost behaviors differed in relation to environmental factors and natural nocturnal moonlight illumination. We predicted that juncos in rural areas would experience less anthropogenic disturbances, such as ALAN, and therefore would initiate roost earlier, depart roost earlier, and have less nocturnal restless periods than urban birds. We also predicted that juncos in rural environments would be more responsive to moonlight compared to urban and suburban birds because cues would be more detectable in rural environments.

## 2. MATERIALS AND METHODS

### 2.1 Study system

Our study was conducted at six sites (2 urban [Hudson and Fairview], 1 suburban [Farhill], 3 rural [Dowler, Woodcock and Baker) in the unglaciated Allegheny Plateau region of southeastern Ohio during the winter of 2025. The rural sites are within a housing intermix embedded in a landscape matrix that is predominantly forested at Dowler and predominantly agricultural (pasture and hay) at Woodcock and Baker. Farhill is a suburban site with woodland and field fragments intermixed with residences and businesses. Fairview and Hudson are urban sites dominated by residential areas and businesses with small patches of trees and shrubs within private property. Streetlamps were common in urban sites and uncommon at rural sites. However, there was a streetlight that illuminated the main roost sites at Dowler (Figure S1). Sites varied in elevation and ALAN (Table S1, Figure 4). Sites were on private land except Woodcock Nature Preserve; however, birds we studied at Woodcock predominately used private land surrounding the site.

**Figure 1.**
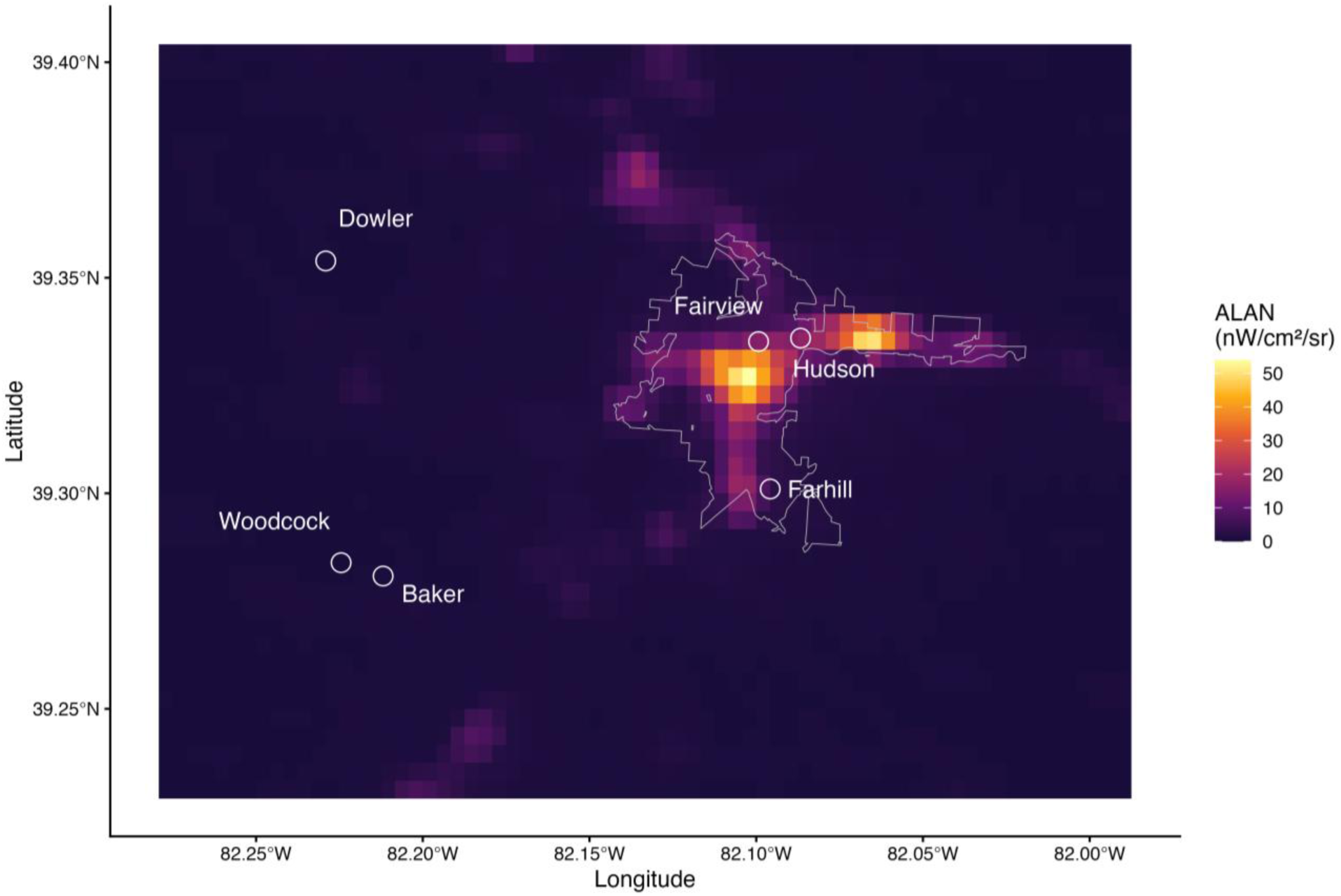
Artificial light at night (ALAN) in Athens County, Ohio. ALAN values (nW/cm²/sr) derived from the VIIRS Black Marble yearly composite product for 2023 (VNP46A4; Román et al. 2018). Grey outline indicates Athens city limits. Circles indicate study sites: Dowler, Woodcock, and Baker (rural) and Fairview, Hudson, and Farhill (within Athens city).

Bird feeders were present on the landscape at all sites and private property owners fed birds during the winter prior to our study. We provided landowners with identical feeder platforms and bird seed mix containing white millet, oil sunflower, safflower, peanuts, and cracked corn (min crude protein = 12%, min crude fat = 21% and max crude fiber = 19%) to ensure similar food availability at all sites and to aid in capture in feeder traps. We captured, banded, and nanotagged (NTQB2-5-1 or NTQB2-2; Lotek Wireless Inc., 166.380 Mhz pulse rates 21.1 – 25.7 seconds) 54 dark-eyed juncos between 2 Jan and 8 Feb 2025.

We used sensorgnome mini-receiving stations with an omnidirectional antenna (166.38 Mhz) mounted on a 3 m pole at Fairview, Hudson, Farhill and Baker. Motus towers (∼ 8.5 m, CTT sensor stations) were already present at Woodcock and Dowler and within Athens (Ohio University, Radar Hill) but the latter did not detect juncos due to topography and distance to the tower. Data were downloaded from Motus for project 406: Ohio Avian Movement Ecology resulting in 14,723,055 detections. We used 7,673,463 detections to examine nocturnal activity patterns. Birds in Athens city were detected on both Hudson and Fairview ministations so data were combined after processing (below) and referred to as Hudson. Birds at Baker and Woodcock were detected on both stations so data were merged during processing and referred to as Baker.

### 2.2 Moonlight, weather and sunrise data

We compiled sunrise, sunset, weather and moonlight data from Jan 1, 2025 – Apr 30, 2025. Sunrise and sunset data were collected for Athens County from https://www.timeanddate.com/sun/@4505546. We used site specific coordinates to gather weather data from the RNCEP package (Kemp et al. 2012) including air temperature (°C), wind speed (m/s), wind direction (degrees), pressure (hPA millibars), atmospheric cloud cover (%) and precipitation rate (kg.m^2^/s). There were few days with precipitation during our study, so this variable was excluded. Weather data were available four times per day, and our study area is in the Eastern time zone (ET), so we assigned UTC = 0 as evening, UTC = 6 as night, UTC = 12 as morning, and UTC **=** 18 as afternoon. We used site specific coordinates to gather moonlight intensity data with the moonlit package (Śmielak 2023, Smielak 2026). We calculated daily moonlight summary statistics for each night by filtering values with function sunAltDegrees < 0 to include only light intensity values when the sun was below the horizon and included mean moonlight illumination (lux) and hours the moon was above the horizon each night. We also deployed HOBO Pendant temperature and light loggers (Model UA-002-64) at each site; however, they were not sensitive enough to record skyglow or luminance at our light polluted sites.

### 2.3 Activity patterns

We used Motus data to estimate activity patterns in the roostR package (Williams 2026). From diel patterns, we calculated roost initiation, roost departure, the proportion of time a bird was detected each night, the number of restless events and the amount of time birds were restless between roost initiation and departure. Because Motus detections are non-continuous bursts of VHF radio waves emitted at user defined pulse rates, the roostR package estimates the number of restless events or proportion restless time per observed hour (Williams 2026). Because the CTT sensor stations had more than one antenna, resulting in more than one detection at a given time, detections were grouped by receiver and antenna. roostR was developed with this data set; all parameters used to extract data for analyses were set at package defaults.

### 2.4 Data analysis

All analyses were conducted in R (v4.6.0; R Core Team 2026). Predictor variables were standardized to provide effect sizes and mean nightly moonlight illumination was log transformed. All variables were back transformed after analyses for biological interpretation. We used individual as a random effect in all models. For each response variable, we constructed a global model based on *a priori* hypotheses and used maximum likelihood (ML) estimation to perform model selection via the dredge function in the MuMIn package (Bartoń 2026), ranking candidate models by AICc. Multicollinearity was assessed using `vif` (package car; Fox and Weisberg 2019)) for linear mixed models or function `check_collinearity` for generalized linear mixed models (package performance; Lüdecke et al. 2021). Selected models were refit with restricted maximum likelihood (REML) for parameter estimation, and residual plots were examined to verify model assumptions. Intraclass correlation coefficients (ICC), marginal R² (variance explained by fixed effects) and conditional R² (variance explained by model) were calculated using the performance package. Post hoc tests with Tukey adjustment for multiple comparisons were performed in emmeans (Lenth and Piaskowski 2026)

#### 2.4.1 Roost initiation and departure

Roost initiation and roost departure were evaluated using mixed effect models in lme4 (Bates et al. 2015). The full model of roost initiation included day of year, site, evening air temperature (°C), mean nightly moonlight illumination (lux), maximum wind speed (m/s), mean cloud cover (%), mean atmospheric pressure (hPa), and the interactions between site and air temperature and site and moonlight illumination. The AICc selected model included all fixed effects except wind and cloud cover (Table S2).

The full model of roost departure included day of year, site, morning air temperature (°C), mean nightly moonlight illumination (lux), maximum wind speed (m/s), mean cloud cover (%), mean atmospheric pressure (hPa), and the interactions between site and air temperature and site and moonlight illumination. The AICc selected model included day of year, site, morning air temperature (°C), mean nightly moonlight illumination (lux), the amount of time the moon was above the horizon, morning wind speed (m/s), overnight cloud cover (%), morning atmospheric pressure (hPa), and the interactions between site and air temperature, site and moonlight illumination (Table S3).

#### 2.4.2 Restless bouts and proportion of time restless

To examine restlessness, we excluded nights where juncos were detected < 40% of the night. Our full model of the number of restless events per hour was used function `glmmTMB` (Brooks et al. 2017) with a gamma distribution and included day of year, site, nighttime air temperature (°C), mean nightly moonlight illumination (lux), nightly wind speed (m/s), the amount of time the moon was above the horizon, nightly cloud cover (%), nightly atmospheric pressure (hPa), and the interactions between site and air temperature and site and moonlight illumination. The AICc selected model included day of year, site, nighttime temperature, mean nightly moonlight illumination, wind speed and cloud cover (Table S4).

Our full model of the proportion of time juncos were restless was modeled using beta regression with a logit link (family = beta; `glmmTMB;` Brooks et al. 2017) and included day of year, site, nighttime air temperature (°C), mean nightly moonlight illumination (lux), nightly wind speed (m/s), the amount of time the moon was above the horizon, nightly cloud cover (%), nightly atmospheric pressure (hPa), and the interactions between site and air temperature and site and moonlight illumination. The AICc selected model included day of year, nighttime air temperature, mean nightly moonlight illumination, site and wind speed (Table S5). Residual plots indicated heterogeneity of dispersion, with juncos at Baker showing the greatest variation in proportion of time restless, so the model was refit with `dispformula = ∼site` to account for site level heterogeneity. We manually calculated conditional and marginal R^2^ using Pearson correlations between the proportion of time restless and fitted and predicted values respectively (Nakagawa and Schielzeth 2013).

## 3. Results

Our data consisted of activity from 54 juncos (Hudson: n = 17, Farhill: n = 9, Baker: n = 13, Dowler: n = 15), totaling 7,215 nights across individuals from 2 January 2025 through 20 April 2025. All juncos but two were second-year birds. Descriptive statistics of roost behaviors and environmental variables by site are in Table S6 and S7.

### 3.1 Roost initiation

Juncos roosted 0.67 ± 0.01 (Baker: mean ± SE) to 1.15 ± 0.01 (Dowler) hours before sunset, roosting later as day length increased but varied by site and in relation to environmental variables. Roost initiation was earlier with increased moonlight illumination, later with warmer temperatures and greater atmospheric pressure (n = 53 juncos on 4582 nights; Table 1, Figure 2). The model explained 71% of the variation in roost initiation with fixed effects accounting for 70% with low individual repeatability (ICC = 0.06). Juncos at Baker roosted later (marginal means: 17.48 ET ± 2.5 min) than all other sites while juncos at Dowler (17:17 ET ± 2.4 min) and Hudson (17:17 ET ± 2.2 min) roosted earliest and Farhill was intermediate (17:17 ET ± 3.1 min; all p < 0.05 except Dowler vs Hudson: p = 0.99). Post hoc tests of the interaction between site and moonlight illumination indicated that birds at Baker showed the strongest negative relationship between roost initiation and moonlight illumination (β = −0.114 ± 0.007 SE, *p* < 0.001, Figure 3), roosting up to 13.8 minutes later on the darkest nights compared to brightest nights, while juncos at Dowler showed the weakest response (β = −0.05 ± 0.007 SE, *p* < 0.001). Hudson and Farhill showed intermediate responses that did not differ from each other (β = −0.088 and −0.057, respectively). While all birds roosted later on warmer evenings, the relationship was not different at Hudson and Farhill (*p* = 0.29) but the response was greater (steeper slopes) at sites within the city limits than at the rural sites (all *p* < 0.001, Figure 4).

**Figure 2.**
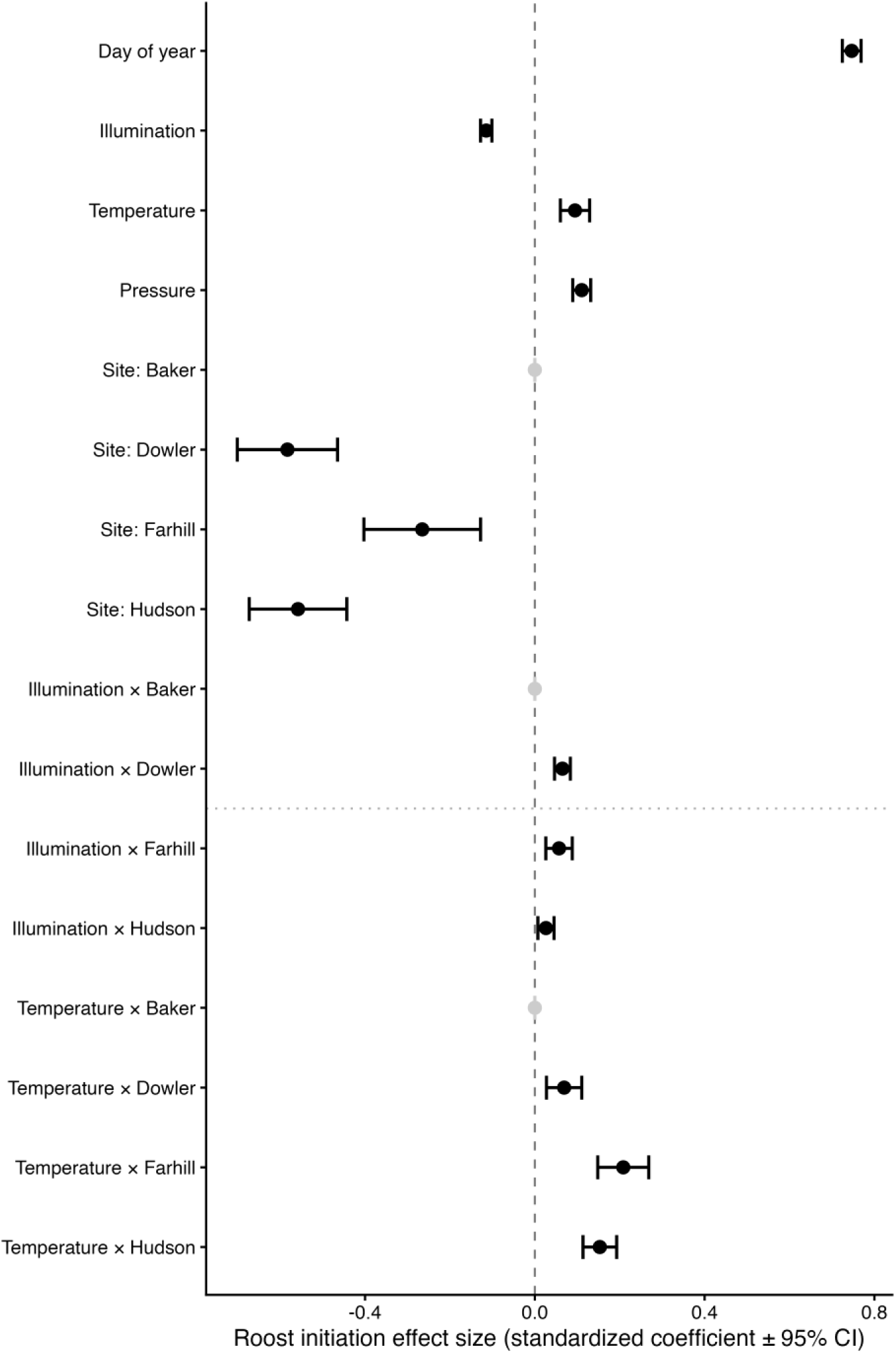
Effect size and standardized coefficients (± 95% CI) of the fixed effects from the model of junco roost initiation.

**Figure 3.**
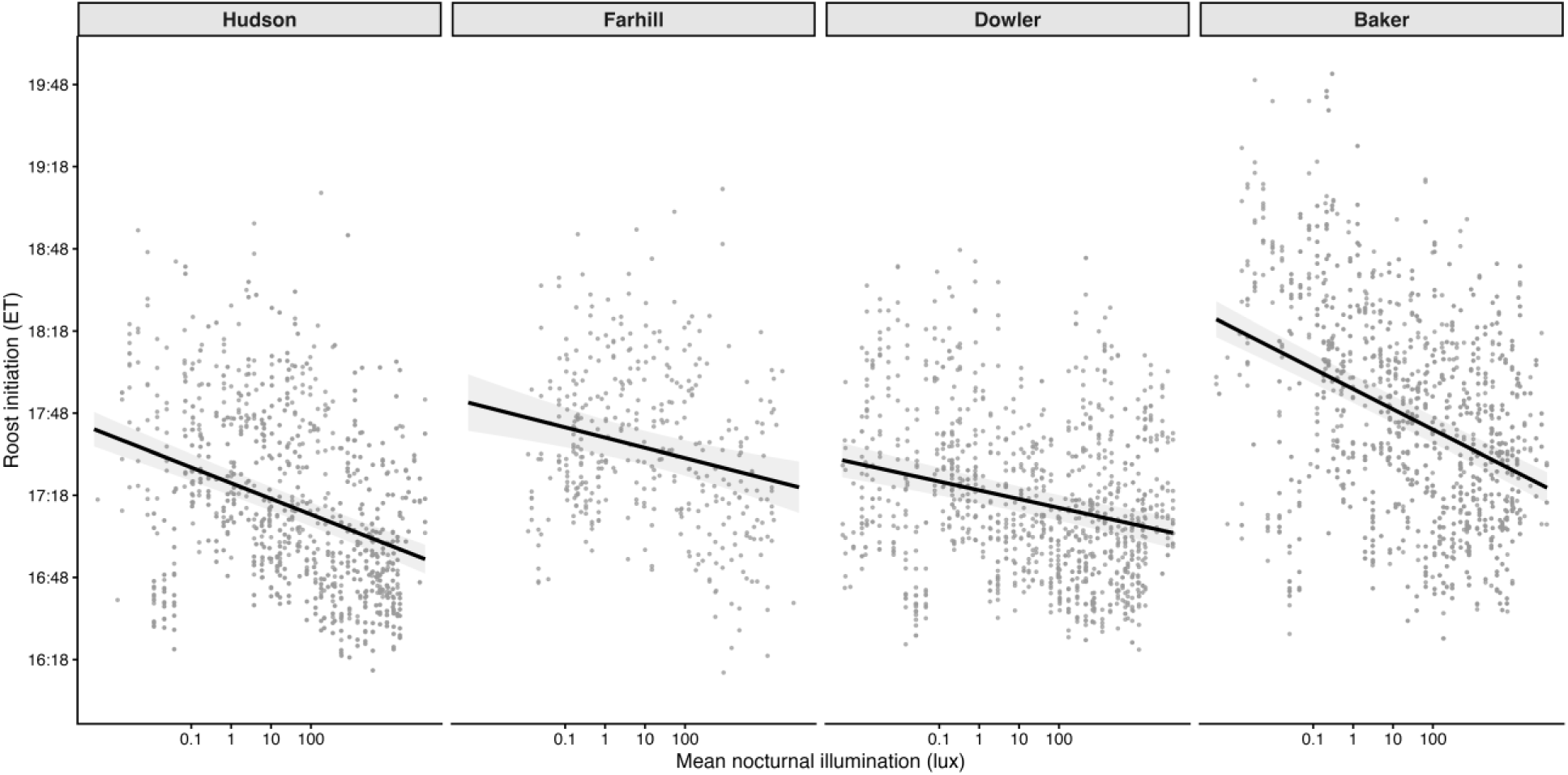
Roost initiation (ET) by juncos in relation to mean moonlight illumination (lux). Moonlight illumination was natural log transformed in the model and back transformed to the response scale for interpretation. Points show partial residuals accounting for other fixed effects in the model and lines show population-level predicted relationships.

**Figure 4.**
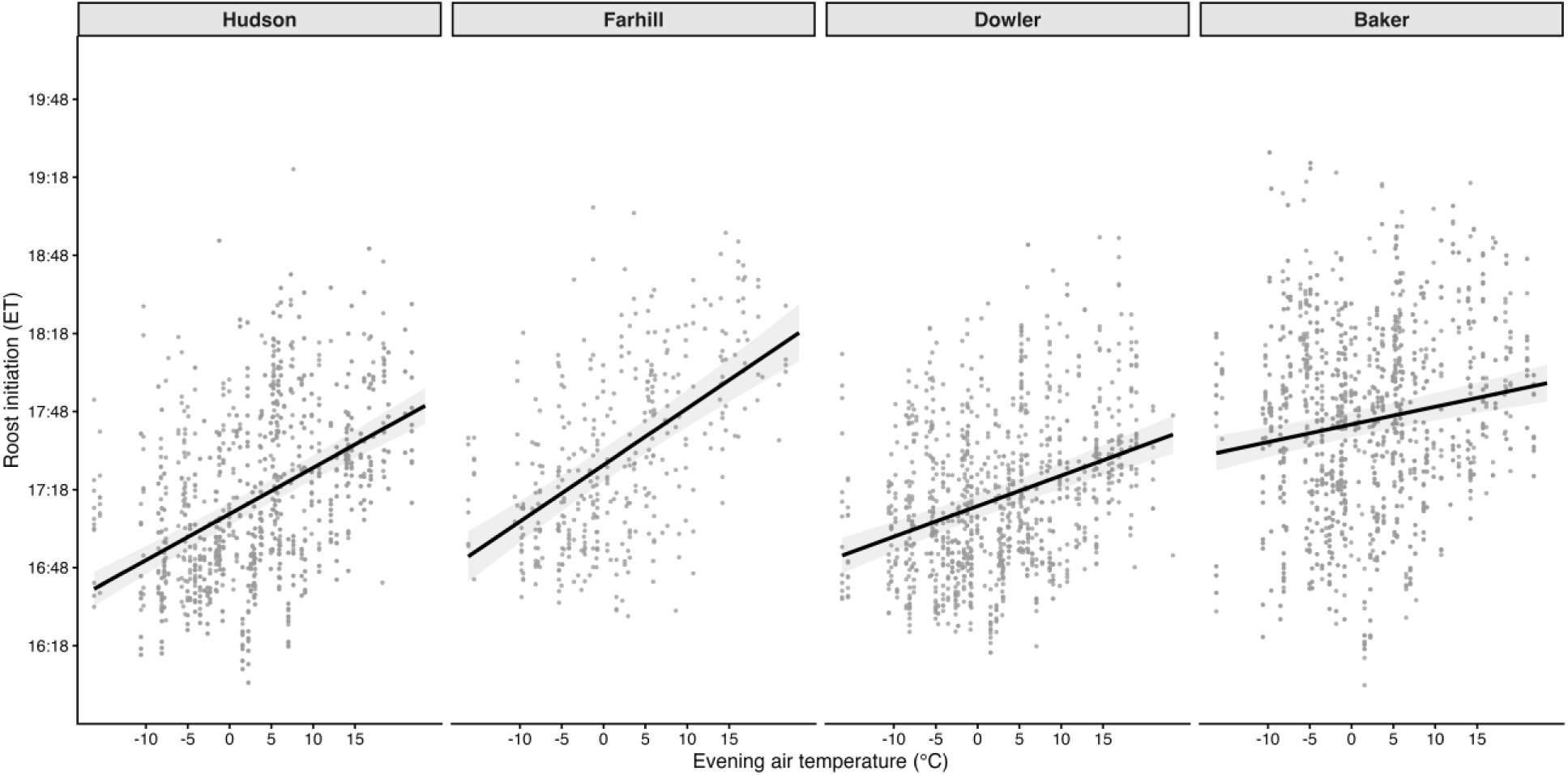
Roost initiation (ET) by juncos in relation to evening air temperature (°C). Points show partial residuals accounting for other fixed effects in the model and lines show population-level predicted relationships.

**Table 1.** Fixed effects from the model of junco roost initiation each evening with individual as a random effect.

| Term | $\beta$ | SE | t | df | p | 95% CI<br>lower | 95% CI<br>upper |
| --- | --- | --- | --- | --- | --- | --- | --- |
| Day of year | 0.747 | 0.011 | 66.251 | 4,566.06 | **0.0000** | 0.724 | 0.769 |
| Site: Dowler | -0.583 | 0.059 | -9.961 | 42.09 | **0.0000** | -0.701 | -0.465 |
| Site: Farhill | -0.265 | 0.068 | -3.879 | 48.81 | **0.0003** | -0.403 | -0.128 |
| Site: Hudson | -0.558 | 0.057 | -9.789 | 42.93 | **0.0000** | -0.673 | -0.443 |
| Temperature | 0.095 | 0.018 | 5.391 | 4,540.99 | **0.0000** | 0.060 | 0.129 |
| Illumination | -0.115 | 0.007 | -16.847 | 4,549.10 | **0.0000** | -0.128 | -0.101 |
| Pressure | 0.110 | 0.011 | 10.225 | 4,527.58 | **0.0000** | 0.089 | 0.132 |
| Temperature × Dowler | 0.069 | 0.021 | 3.256 | 4,552.2 | **0.0011** | 0.027 | 0.111 |
| Temperature × Farhill | 0.208 | 0.031 | 6.799 | 4,566.93 | **0.0000** | 0.148 | 0.268 |
| Temperature × Hudson | 0.153 | 0.020 | 7.571 | 4,567.48 | **0.0000** | 0.113 | 0.193 |
| Illumination × Dowler | 0.065 | 0.010 | 6.841 | 4,566.99 | **0.0000** | 0.046 | 0.084 |
| Illumination × Farhill | 0.057 | 0.016 | 3.588 | 4,566.33 | **0.0003** | 0.026 | 0.088 |
| Illumination × Hudson | 0.026 | 0.010 | 2.697 | 4,564.31 | **0.0070** | 0.007 | 0.045 |

$\beta$ = standardized coefficient; SE = standard error; CI = confidence
interval. Bold p-values indicate significance at $\alpha = 0.05$ .

### 3.2 Roost Departure

Juncos departed roosts 0.26 ± 0.02 hours before sunrise (Baker) to 0.03 ± 0.02 hours after sunrise (Hudson), departing roosts earlier as day length increased but varied by site and in relation to environmental variables (n = 53 juncos on 6286 nights; Table 2, Figure 5). Juncos departed earlier with increased moonlight illumination, wind speed and cloud cover but later in relation to warmer temperatures, the amount of time the moon was above the horizon and atmospheric pressure. The model explained 32% of the variation in roost departure with fixed effects accounting for 26% and indicated low individual repeatability (ICC = 0.08). Juncos at Baker departed roosts consistently earlier (marginal mean: 7:10 ET ± 2.1 min) than all other sites (p < 0.001), while juncos at Dowler (7.19 ET ± 2 min) and Farhill (7.35 ± 0.04) departed roosts similarly (p = 0.92), and juncos at Hudson departed the latest (7:28 ET ± 1.9 min) in relation to juncos at Baker and Dowler (p < 0.01) but no difference compared to Farhill (p = 0.08). Post hoc tests of the interaction indicated that juncos at Farhill had the strongest response to moonlight illumination compared to all other sites (β = −0.081 ± 0.008, *p* < 0.001, Figure 6), departing 9.6 minutes earlier after brighter nights. Juncos at Baker and Dowler responded similarly (p = 0.99) with the weakest response, while juncos at Hudson showed an intermediate response to moonlight illumination (β = −0.038 ± 0.005, *p* < 0.001). Juncos departed later on warmer mornings (*p* < 0.001; Figure 7) with juncos at Hudson showing the strongest response (β = 0.231 ± 0.013, *p* < 0.001), departing approximately 27.6 minutes later on warm mornings compared to cold mornings. Juncos at Farhill showed the weakest response to temperature (β = 0.059 ± 0.018, *p* = 0.001), while juncos at Baker and Dowler showed similar intermediate responses (Baker: β = 0.176 ± 0.012; Dowler: β = 0.179 ± 0.013, both *p* < 0.001; Baker vs Dowler *p* = 0.99 at all temperature levels).

**Figure 5.**
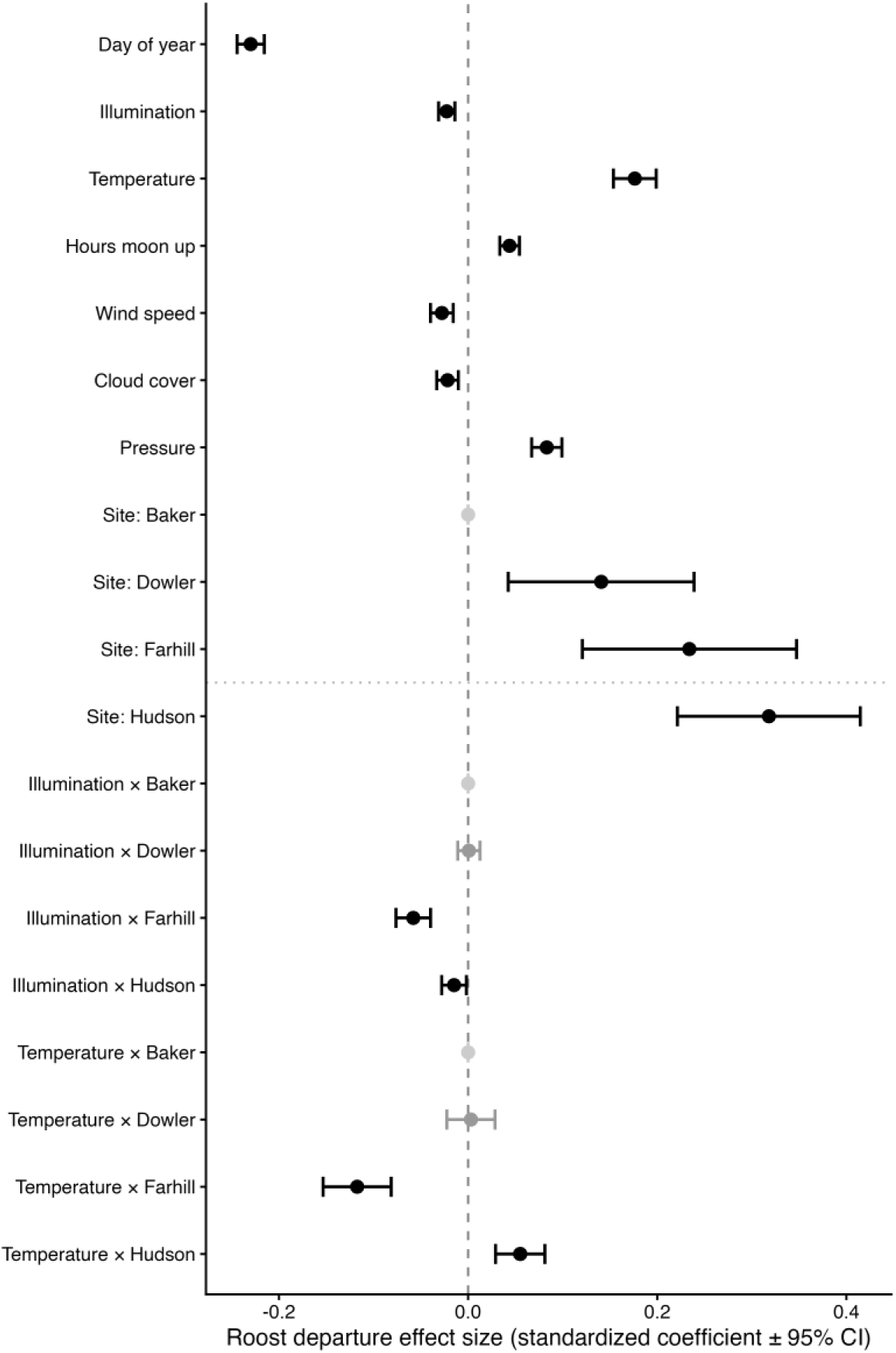
Effect size and standardized coefficients (± 95% CI) of the fixed effects from the model of junco roost departure.

**Figure 6.**
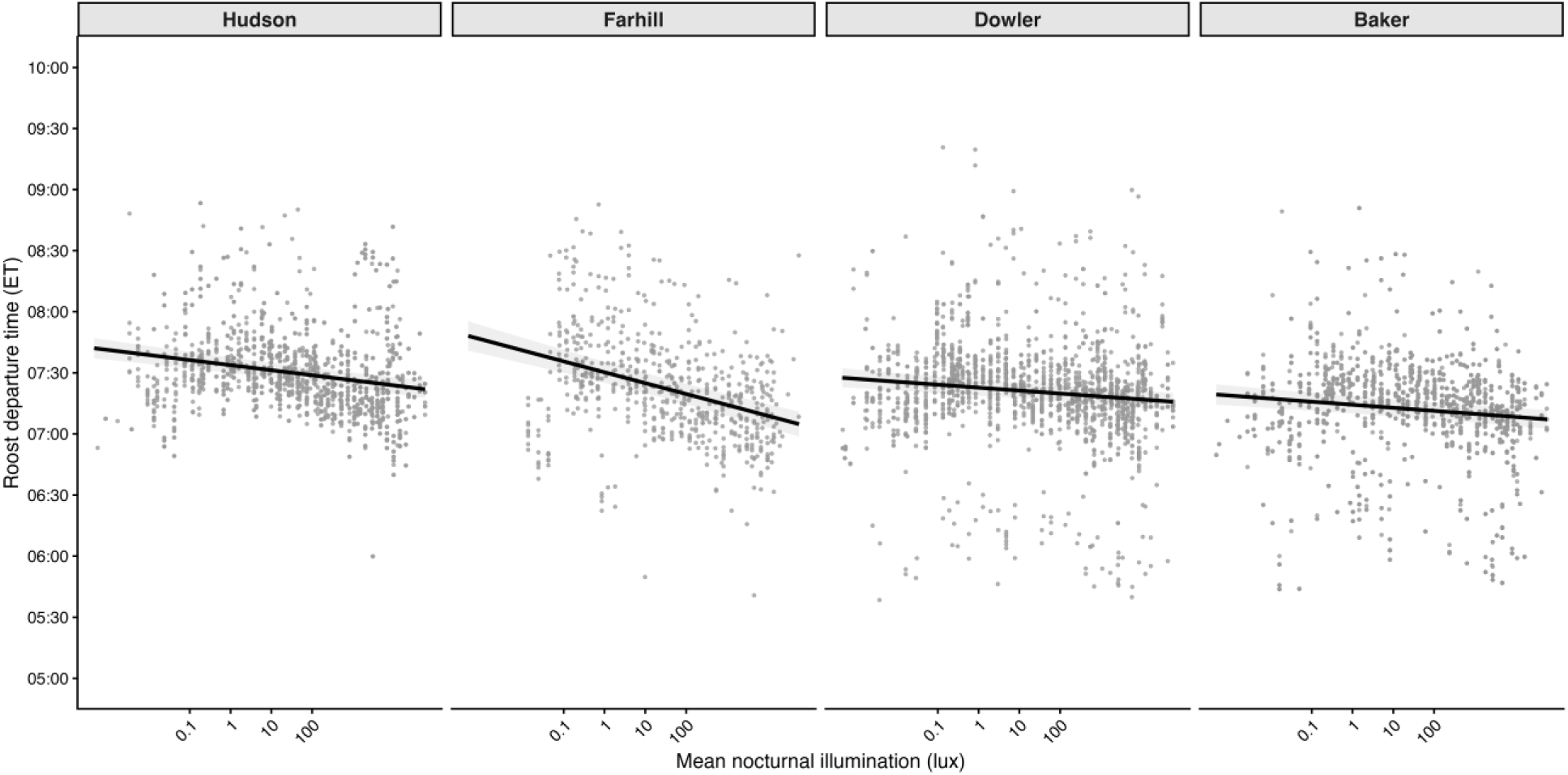
Roost departure (ET) of juncos in relation to mean moonlight illumination (lux). Moonlight illumination was natural log transformed in the model and back transformed to the response scale for interpretation. Points show partial residuals accounting for other fixed effects and lines show population-level predicted relationships.

**Figure 7.**
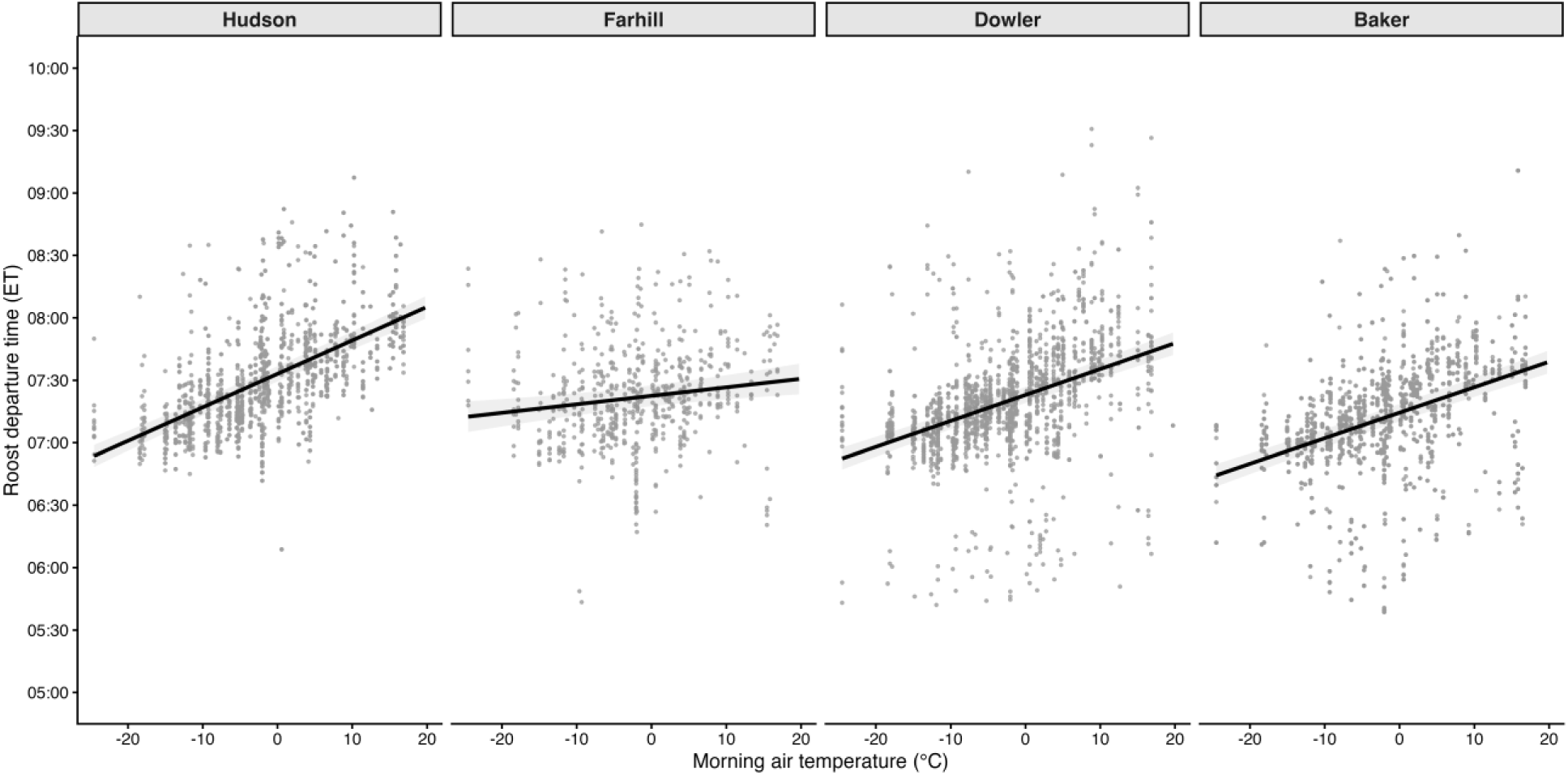
Roost departure (ET) of juncos in relation to morning air temperature (°C). Points show partial residuals accounting for other fixed effects and lines show population-level predicted relationhsips.

**Table 2.**
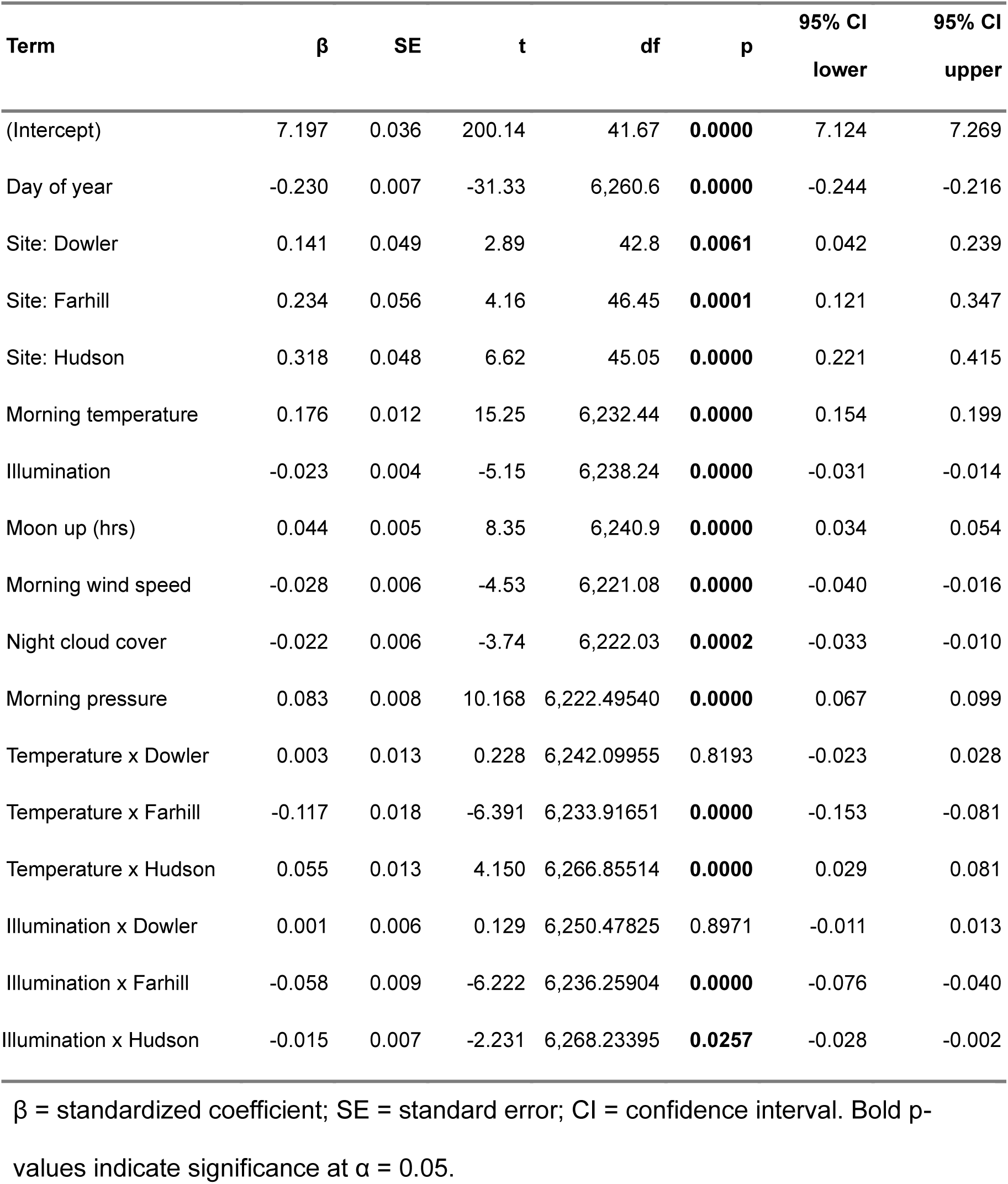
Fixed effects from the model of junco roost departure each morning with individual as a random effect.

| Term | $\beta$ | SE | t | df | p | 95% CI<br>lower | 95% CI<br>upper |
| --- | --- | --- | --- | --- | --- | --- | --- |
| (Intercept) | 7.197 | 0.036 | 200.14 | 41.67 | **0.0000** | 7.124 | 7.269 |
| Day of year | -0.230 | 0.007 | -31.33 | 6,260.6 | **0.0000** | -0.244 | -0.216 |
| Site: Dowler | 0.141 | 0.049 | 2.89 | 42.8 | **0.0061** | 0.042 | 0.239 |
| Site: Farhill | 0.234 | 0.056 | 4.16 | 46.45 | **0.0001** | 0.121 | 0.347 |
| Site: Hudson | 0.318 | 0.048 | 6.62 | 45.05 | **0.0000** | 0.221 | 0.415 |
| Morning temperature | 0.176 | 0.012 | 15.25 | 6,232.44 | **0.0000** | 0.154 | 0.199 |
| Illumination | -0.023 | 0.004 | -5.15 | 6,238.24 | **0.0000** | -0.031 | -0.014 |
| Moon up (hrs) | 0.044 | 0.005 | 8.35 | 6,240.9 | **0.0000** | 0.034 | 0.054 |
| Morning wind speed | -0.028 | 0.006 | -4.53 | 6,221.08 | **0.0000** | -0.040 | -0.016 |
| Night cloud cover | -0.022 | 0.006 | -3.74 | 6,222.03 | **0.0002** | -0.033 | -0.010 |
| Morning pressure | 0.083 | 0.008 | 10.168 | 6,222.49540 | **0.0000** | 0.067 | 0.099 |
| Temperature x Dowler | 0.003 | 0.013 | 0.228 | 6,242.09955 | 0.8193 | -0.023 | 0.028 |
| Temperature x Farhill | -0.117 | 0.018 | -6.391 | 6,233.91651 | **0.0000** | -0.153 | -0.081 |
| Temperature x Hudson | 0.055 | 0.013 | 4.150 | 6,266.85514 | **0.0000** | 0.029 | 0.081 |
| Illumination x Dowler | 0.001 | 0.006 | 0.129 | 6,250.47825 | 0.8971 | -0.011 | 0.013 |
| Illumination x Farhill | -0.058 | 0.009 | -6.222 | 6,236.25904 | **0.0000** | -0.076 | -0.040 |
| Illumination x Hudson | -0.015 | 0.007 | -2.231 | 6,268.23395 | **0.0257** | -0.028 | -0.002 |

$\beta$ = standardized coefficient; SE = standard error; CI = confidence interval. Bold p-values indicate significance at $\alpha = 0.05$ .

### 3.3 Restless bouts and proportion of time restless

Juncos displayed from 0.83 ± 0.04 (Hudson) to 2.79 ± 0.19 (Baker) restless events per hour, increasing the number of restless events with warmer temperatures, moonlight illumination, greater wind speed and cloud cover (n = 52 individuals, 2268 nights; Table 3, Figure 8). The model explained 50% of the variation in restless bouts with fixed effects accounting for 39% and some individual repeatability (ICC = 0.18). Juncos at Baker had the most restless bouts (marginal mean: 2.74 ± 0.26 events per hour) compared to all other sites (all *p* < 0.001) followed by juncos at Farhill (1.72 ± 0.20 events per hour, all contrasts *p* < 0.01), while there were no differences in the number of restless events between Dowler (1.11 ± 0.10) and Hudson (0.80 ± 0.07; *p* = 0.31).

**Figure 8.**
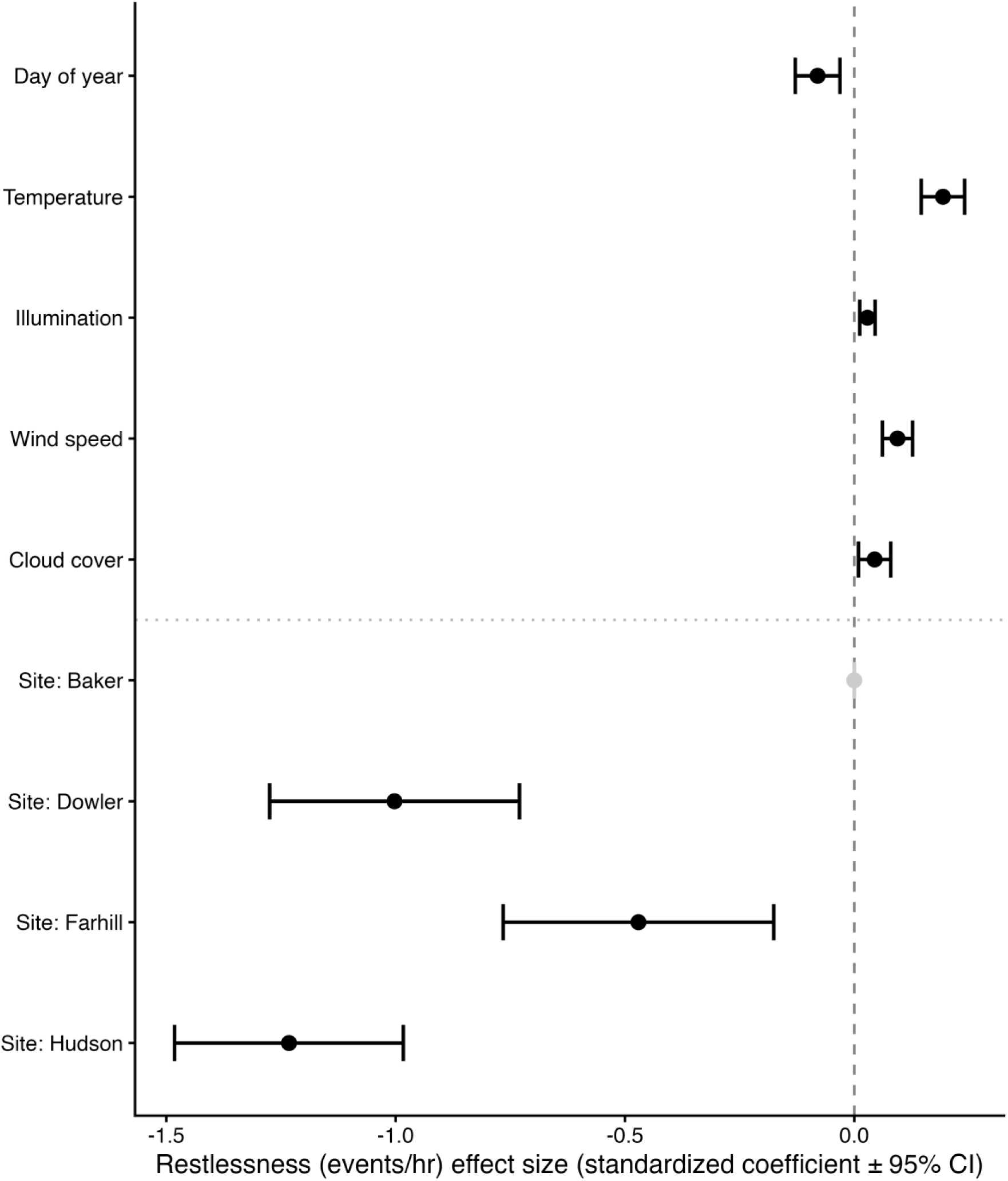
Effect size and standardized coefficients (± 95% CI) of the fixed effects from the model of junco restlessness (events/hr).

**Table 3.**
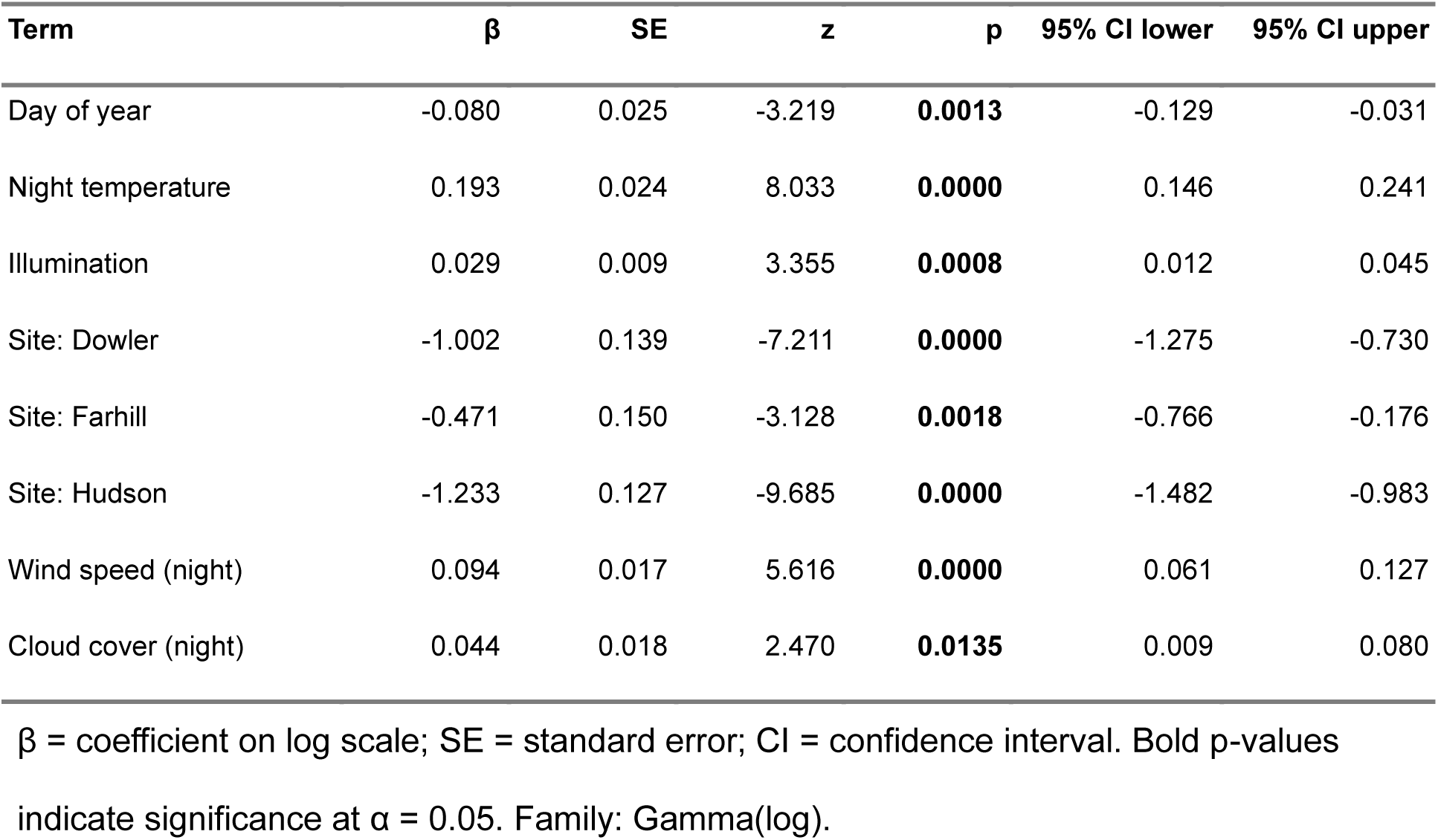
Fixed effects from the model of junco restless rate (events/hr) with individual as a random effect.

| Term | $\beta$ | SE | z | p | 95% CI lower | 95% CI upper |
| --- | --- | --- | --- | --- | --- | --- |
| Day of year | -0.080 | 0.025 | -3.219 | **0.0013** | -0.129 | -0.031 |
| Night temperature | 0.193 | 0.024 | 8.033 | **0.0000** | 0.146 | 0.241 |
| Illumination | 0.029 | 0.009 | 3.355 | **0.0008** | 0.012 | 0.045 |
| Site: Dowler | -1.002 | 0.139 | -7.211 | **0.0000** | -1.275 | -0.730 |
| Site: Farhill | -0.471 | 0.150 | -3.128 | **0.0018** | -0.766 | -0.176 |
| Site: Hudson | -1.233 | 0.127 | -9.685 | **0.0000** | -1.482 | -0.983 |
| Wind speed (night) | 0.094 | 0.017 | 5.616 | **0.0000** | 0.061 | 0.127 |
| Cloud cover (night) | 0.044 | 0.018 | 2.470 | **0.0135** | 0.009 | 0.080 |

$\beta$ = coefficient on log scale; SE = standard error; CI = confidence interval. Bold p-values indicate significance at $\alpha = 0.05$ . Family: Gamma(log).

The proportion of time restless increased with nightly temperature and wind speed but did not differ with day of year or moonlight illumination (Table 4, Figure 9). Fixed and random effects explained 23.3% of variance in the proportion of time restless while the environmental predictors explained 18.5% and some individual repeatability (ICC = 0.156). Juncos at Baker spent the greatest proportion of roosting time restless (marginal mean: 2.28% ± 0.14%), being 2.5 times more restless than juncos at Dowler (odds ratio [OR] = 2.51, *p* < 0.001), 1.7 times more restless than Farhill (OR = 1.67, *p* < 0.001), and 3.6 times more restless than Hudson (OR = 3.55, *p* < 0.001). Juncos at Farhill spent an intermediate proportion of time restless (1.38% ± 0.12%), differing significantly from all other sites (all *p* < 0.003). Juncos at Dowler (0.92% ± 0.07% SE) and Hudson (0.65% ± 0.04% SE) spent the least time restless, though these sites also differed significantly from each other (OR = 1.42, *p* = 0.002)

**Figure 9.**
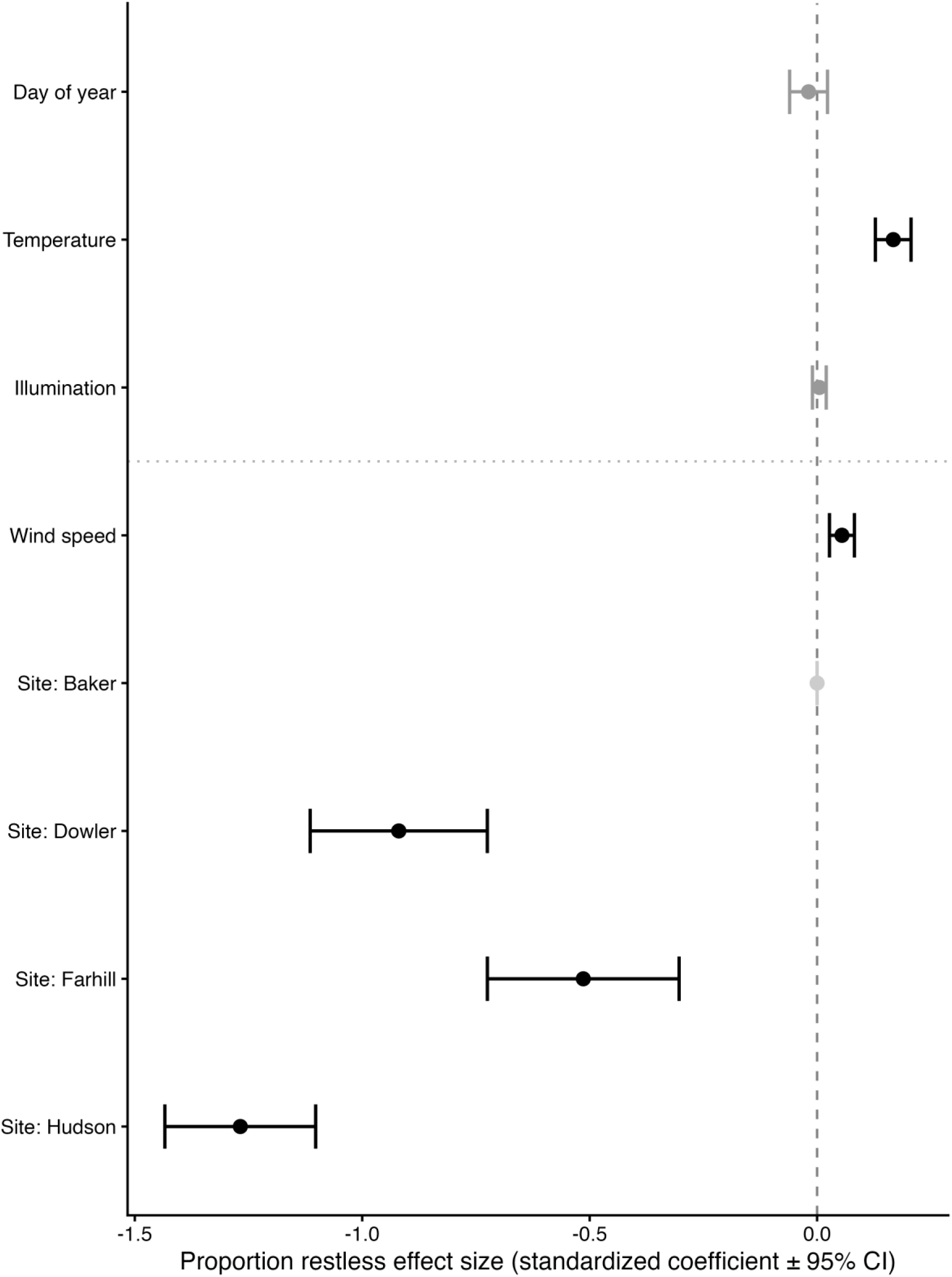
Effect size and standardized coefficients (± 95% CI) of the fixed effects from the model of the proportion of time juncos were restless.

**Table 4.**
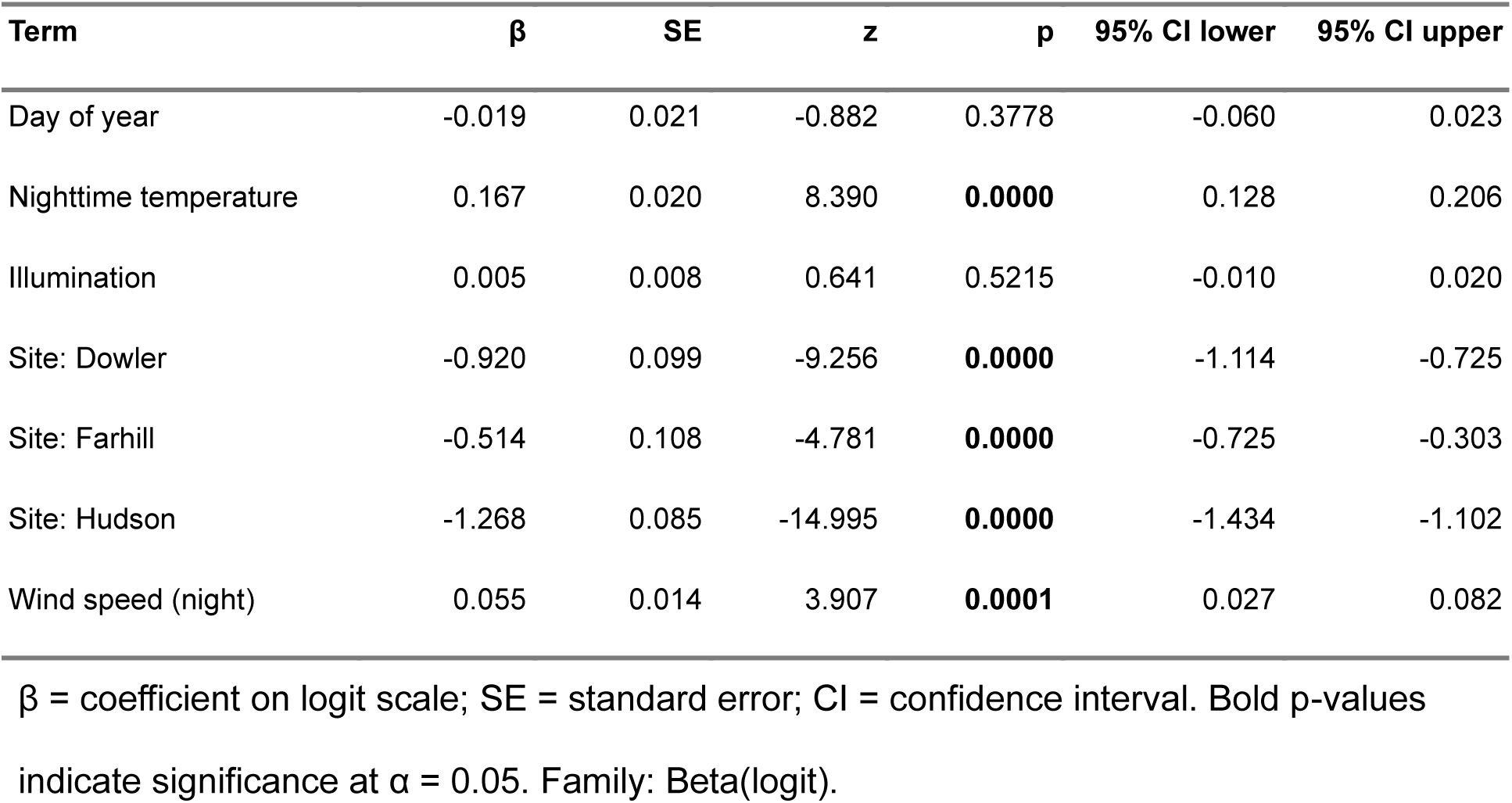
Fixed effects from the model of the proportion of time juncos were restless each night with individual as a random effect and variance structure to account for heterogeneity of variance by site.

## 4. Discussion

Overwintering passerines in temperate zones experience a variety of environmental constraints, which intensify at night, when temperatures fall and foraging is reduced. Overwintering birds may enhance survival by adjusting nocturnal behavior in response to environmental cues (Dobbs and Martin 2000). However, anthropogenic factors can alter the strength, detectability, and importance of natural cues, which can affect how responsive individuals are to different cues (Russart and Nelson 2018, Dickerson et al. 2022, Preston and Brigham 2023). We found that nocturnal behavior of overwintering juncos was modulated by environmental factors, including moonlight illumination and temperature. Birds use light cues and temperature to regulate nocturnal behaviors (Lehmann et al. 2012, Rubolini et al. 2015, Bianco et al. 2025, Drewitt et al. 2026). However, the strength of the response to moonlight illumination and temperature differed among urban, suburban, and rural sites, suggesting that natural cues of moonlight were masked in urban areas and in a rural site, Dowler, where a streetlight was present near roost sites. The interactions between site, temperature, and moonlight illumination support our prediction that local environmental conditions, such as ALAN and urbanization, affect the responsiveness to different environmental cues that are used to modulate nocturnal behavior.

We also predicted that urban and suburban juncos would roost later, rise earlier, and be more restless than rural birds. However, our results did not support these predictions, possibly due to interactions between environmental conditions and resource availability. Although we did not formally analyze roost availability, available data suggest that Dowler and Hudson had lower coniferous cover than Baker and Farhill (Morgan and Williams, in prep). Juncos use coniferous tress for roosting because dense conifer cover offers greater insulation and reduced heat loss from wind exposure (Gottfried and Franks 1975, Chandler et al. 1995). In many species, individuals that arrive later to a limited, high-quality roost are limited to lower-quality positions (Nomano et al. 2021, Lowney and Thomson 2026). Similarly, Allen (2025) found that dark-eyed juncos arrived at roost sites up to 2 hours before sunset to secure thermally beneficial sites.

### 4.1 Roost Initiation

We expected birds would delay roosting on bright nights (e.g., Alonso et al. 1985), but juncos initiated roost earlier on nights with greater moonlight illumination and the strength of the relationship varied by site. Juncos at Baker (rural) initiated roost latest and had the strongest relationship with moonlight illumination, whereas juncos at Farhill (suburban) and Hudson (urban) were less sensitive. While Dowler was classified as a dark site, roost initiation at Dowler had the weakest relationship with moonlight illumination. Our findings may be explained by the mismatch between landscape-level light classifications and local light conditions and roost availability. Juncos at Dowler roosted within a small patch of coniferous trees exposed to light trespass from a streetlamp. A close-range light source can disproportionately shape behavior even when it contributes little to landscape-scale artificial illumination (Van Doren et al. 2017, Adams et al. 2024). By contrast, the birds at Baker tended to roost away from anthropogenic light sources, often within a large pine stand. The strong association between higher levels of moonlight illumination and later roost initiation at Baker may have reflected the birds’ ability to track light cues and make trade-offs between foraging and roosting behaviors (Rubolini et al. 2015, Bianco et al. 2025, Drewitt et al. 2026).

Birds initiated roosting later under more warmer temperatures, but the relationship between initiation and temperature was stronger at urban and suburban sites. Temperature is often used in conjunction with illumination as a signal for roost initiation (Reebs 1986, Schlicht and Kempenaers 2020) because thermal conditions directly reflect energetic constraints birds face in winter (Swanson et al. 2014, Li et al. 2022). When ALAN masks moon illumination cues, alternative cues may be relied on more. As such, juncos overwintering at urban sites may have been more responsive to temperature because temperature was a more reliable signal.

### 4.2 Restlessness

Restlessness varied with environmental conditions and among sites, with juncos at Baker more restless than those at urbanized sites. The number of nocturnal restless bouts increased with warmer temperatures, greater moonlight illumination, wind speed and cloud cover, and the proportion of time spent restless each night increased with temperature and wind speed. While moonlit conditions can elevate visibility for nocturnal predators and increase the perceived predation risks for birds that remain active during the evening (Schmidt 2006, Jolkkonen et al. 2023, Taylor et al. 2023), moonlight can act as a behavioral stimulant for nocturnal activity, increasing foraging, vocalization, and display behavior (Pérez-Granados and Traba 2021, Buda et al. 2025, Alonso et al. 2026). In contrast, prior work found nocturnal activity is typically buffered against temperature variation (Caorsi et al. 2019, Pérez-Granados and Traba 2021) and suppressed by wind (Prinz et al. 2025, Diaz-Palma et al. 2026).

Restlessness also differed among sites. Juncos at Baker, and to a lesser extent, Farhill, were more restless than Dowler and Hudson. These results may be explained by ALAN and/or roost availability. At Dowler birds roosted within light trespass of a streetlamp and Hudson is urban with high ALAN. ALAN increases the visibility and detectability of prey species for predators (Negro et al. 2020, Rodríguez et al. 2021, Adams et al. 2024) and birds often remain still in the presence of predators (Mouritsen 1992, Bianco et al. 2025). Mesopredator populations reach higher densities in residential developments and wildland-urban intermix zones, driven by anthropogenic food subsidies and declines of apex predators (Rodríguez et al. 2021, Veon et al. 2023, Leighton et al. 2023), and urban areas host higher numbers of outdoor cats (Rodewald et al. 2011). Juncos roosting in artificially lit, roost limited sites may reduce movement and restlessness to lower predation risk and avoid losing the roost position to a competitor.

### 4.3 Roost Departure

Across sites, juncos departed earlier on nights with greater moonlight illumination, possibly due to increased wakefulness, or alertness (Kronfeld-Schor et al. 2013, Evens et al. 2020). In contrast, juncos departed later after warmer nights consistent with Williams et al. (2020, 2025) where birds initiated morning activity earlier on cold conditions, likely because greater overnight energy expenditure from lower temperatures prompt earlier foraging (Reid et al. 2000, Conway 2000, Conway and Martin 2000).

Roost departure also differed by site: juncos at Baker departed the earliest, followed by Farhill and Dowler, then Hudson. Juncos at Hudson showed the strongest response to temperature, likely because urban heat islands buffer birds from the thermoregulatory costs relative to colder, rural sites (Yang and Bou-Zeid 2018, Bonnet-Lebrun et al. 2020, Sumasgutner et al. 2023). Differences in restlessness among the sites may partially explain these findings. Restlessness carries an energetic cost (Ouyang et al. 2017, Ferretti et al. 2019, Litt-Jukes and Morrissey 2026), and birds often adjust behavior when reserves are limited (Ferretti et al. 2019, Barratt et al. 2025). Because juncos at Baker and Farhill were more restless, these birds likely incurred greater overnight energy expenditures and departed roosts earlier to replenish depleted reserves. Rural areas also tend to offer fewer reliable food resources than urban areas, which have higher densities of bird feeders, food waste, and other anthropogenic subsidies (Tryjanowski et al. 2015). Together, higher thermoregulatory costs, restlessness-driven energy expenditure, and food availability may help explain why juncos at Baker, Dowler, and Farhill departed roosts earlier than at Hudson.

## 5 Conclusions

Lunar cycling shapes activity, sleep, and behavior across many species (Yamamoto and Trathan 2015, Portugal et al. 2019, Van Hasselt et al. 2020) and ALAN alters activity patterns (Dominoni et al. 2013, Van Doren et al. 2017, Diaz-Palma et al. 2026, Champenois et al. 2026). Our results suggest that ALAN can mask the natural light cues birds use to time roosting, prompting a shift toward temperature as an alternative cue, and representing a key mechanism by which human-altered landscapes can disrupt nocturnal ecology in birds. Roost behaviors in juncos were further shaped by weather. The availability of thermally favorable roost sites and pressures from predators or competitors may also have contributed, though we did not directly measure these factors; these relationships warrant further study. On altered landscapes, ALAN likely interacts with environmental conditions, roost availability, and predation or competition pressures to amplify signal disruption and drive behavioral modifications. Mitigation measures to reduce light pollution can benefit human and animal health (Svechkina et al. 2020, Yang et al. 2024, Thomas et al. 2025) and minimize disruption to diel activity patterns in overwintering birds.

