## Supplemental figures and tables for "Overwinter roost activity patterns in response to environmental conditions along a rural-urban gradient"

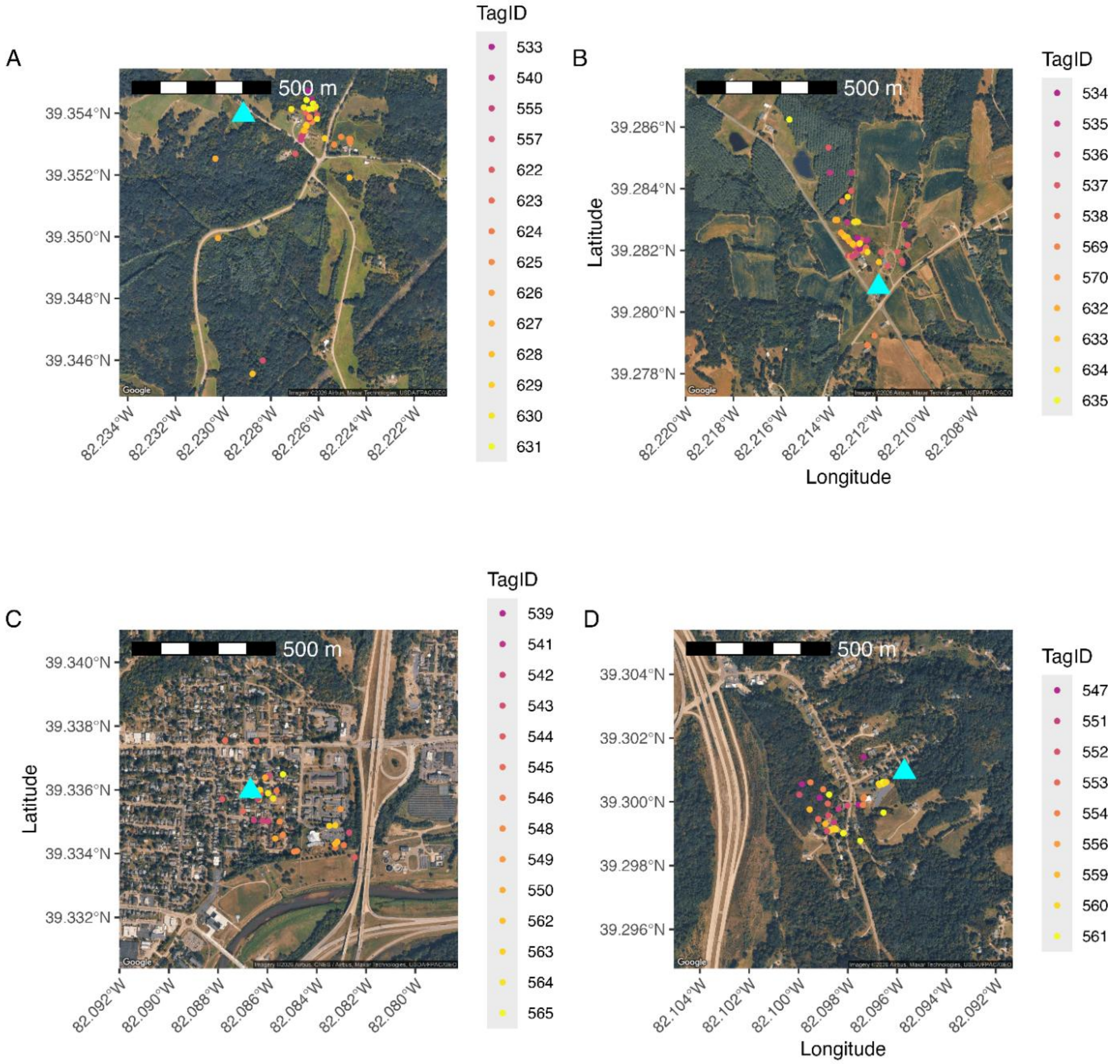

Figure S1. Junco roost site detections during manual radio telemetry at A) Dowler, B) Baker, C) Hudson, and D) Farhills. Colored points represent different juncos (TagID), and blue triangles represent the location of a Motus receiving station. At Dowler, there was a streetlight across the street from the main cluster of roost locations.

Table S 1 Study site locations, coordinates, elevation and ALAN.

| Site | Latitude | Longitude | Elevation (m) | ALAN (nW/cm²/sr) |
|---|---|---|---|---|
| Farhill | 39.3010 | -82.0958 | 231.7 | 2.75 |
| Hudson | 39.3360 | -82.0867 | 193.9 | 17.26 |
| Fairview | 39.3352 | -82.0993 | 256.3 | 15.59 |
| Baker | 39.2808 | -82.2119 | 258.6 | 0.58 |
| Woodcock | 39.2839 | -82.2245 | 282.1 | 0.00 |
| Dowler | 39.3539 | -82.2291 | 288.7 | 0.00 |

ALAN = artificial light at night derived from VIIRS Black Marble annual composite (VNP46A4, 2023; Román et al. 2018). Elevation derived from USGS Elevation Point Query Service.

Table S 2 Model selection table for the fixed effects of roost initiation (time) by juncos including models with ΔAICc < 10.

| (Intercept) | Temp | Cloud | DoY | logIllum | Pres | Site | Wind | Temp × Site | logIllum × Site | K | logLik | AICc | ΔAICc | w |
|---|---|---|---|---|---|---|---|---|---|---|---|---|---|---|
| **17.91645** | **+** | | **+** | **+** | **+** | **+** | | **+** | **+** | **16** | **-3,553.64** | **7,139.41** | **0.00** | **0.42** |
| 17.91565 | + | | + | + | + | + | + | + | + | 17 | -3,552.99 | 7,140.12 | 0.71 | 0.29 |
| 17.91650 | + | + | + | + | + | + | | + | + | 17 | -3,553.54 | 7,141.21 | 1.80 | 0.17 |
| 17.91570 | + | + | + | + | + | + | + | + | + | 18 | -3,552.88 | 7,141.92 | 2.51 | 0.12 |

Table S 3 Model selection table for the fixed effects of roost departure (time) by juncos including models with ΔAICc < 10.

| (Intercept) | Temp | Cloud | DoY | HrsMoon | logIllum | Precip | Pres | Site | Wind | Temp × Site | logIllum x Hrs | logIllum × Site | K | logLik | AICc | ΔAICc | w |
|---|---|---|---|---|---|---|---|---|---|---|---|---|---|---|---|---|---|
| **7.196223** | **+** | **+** | **+** | **+** | **+** | | **+** | **+** | **+** | **+** | | **+** | **19** | **-3,084.38** | **6,206.89** | **0.00** | **0.38** |
| 7.200166 | + | + | + | + | + | | + | + | + | + | + | + | 20 | -3,083.49 | 6,207.11 | 0.22 | 0.34 |
| 7.196338 | + | + | + | + | + | + | + | + | + | + | | + | 20 | -3,084.35 | 6,208.84 | 1.95 | 0.14 |
| 7.200257 | + | + | + | + | + | + | + | + | + | + | + | + | 21 | -3,083.46 | 6,209.07 | 2.18 | 0.13 |

Table S 4 Model selection table for the fixed effects of restless events per hour by juncos including models with ΔAICc < 10.

| Temp | Cloud | DoY | Moon hrs | Illum | Pres | Site | Wind | Illum × Site | K | logLik | AICc | ΔAICc | w |
|---|---|---|---|---|---|---|---|---|---|---|---|---|---|
| **+** | **+** | **+** | | **+** | | **+** | **+** | | **11** | **-2,937.49** | **5,897.09** | **0.00** | **0.29** |
| + | + | + | + | + | | + | + | | 12 | -2,936.98 | 5,898.10 | 1.01 | 0.18 |
| + | + | + | | + | + | + | + | | 12 | -2,937.16 | 5,898.47 | 1.38 | 0.15 |
| + | + | + | + | + | + | + | + | | 13 | -2,936.54 | 5,899.24 | 2.15 | 0.10 |
| + | + | + | | + | | + | + | + | 14 | -2,936.39 | 5,900.98 | 3.89 | 0.04 |
| + | | + | + | + | | + | + | | 11 | -2,939.44 | 5,901.00 | 3.91 | 0.04 |
| + | | + | | + | | + | + | | 10 | -2,940.61 | 5,901.32 | 4.23 | 0.04 |
| + | | + | + | + | + | + | + | | 12 | -2,938.69 | 5,901.53 | 4.44 | 0.03 |
| + | + | + | + | + | | + | + | + | 15 | -2,935.79 | 5,901.80 | 4.71 | 0.03 |
| + | | + | | + | + | + | + | | 11 | -2,940.04 | 5,902.19 | 5.10 | 0.02 |
| + | + | + | | + | + | + | + | + | 15 | -2,936.05 | 5,902.31 | 5.22 | 0.02 |
| + | + | + | + | + | + | + | + | + | 16 | -2,935.31 | 5,902.86 | 5.78 | 0.02 |
| + | | + | + | + | | + | + | + | 14 | -2,938.13 | 5,904.45 | 7.36 | 0.01 |
| + | | + | + | + | + | + | + | + | 15 | -2,937.34 | 5,904.89 | 7.80 | 0.01 |
| + | | + | | + | | + | + | + | 13 | -2,939.45 | 5,905.06 | 7.97 | 0.01 |
| + | + | | | + | + | + | + | | 11 | -2,941.57 | 5,905.25 | 8.16 | 0.00 |
| + | + | | | + | | + | + | | 10 | -2,942.63 | 5,905.36 | 8.27 | 0.00 |
| + | | + | | + | + | + | + | + | 14 | -2,938.84 | 5,905.87 | 8.78 | 0.00 |
| + | + | | + | + | + | + | + | | 12 | -2,940.90 | 5,905.94 | 8.85 | 0.00 |
| + | + | + | | | | + | + | | 10 | -2,943.07 | 5,906.24 | 9.15 | 0.00 |
| + | + | | + | + | | + | + | | 11 | -2,942.18 | 5,906.48 | 9.39 | 0.00 |

Table S 5 Model selection table for the fixed effects of the proportion of time juncos were restless including models with ΔAICc < 2.

| Temp | Cloud | DoY | Moon hrs | Illum | Pres | Site | Wind | IllumxCloud | IllumxMoon | Illum×Site | K | logLik | AICc | ΔAICc | w |
|---|---|---|---|---|---|---|---|---|---|---|---|---|---|---|---|
| **+** | | **+** | | **+** | | **+** | **+** | | | | **10** | **7,715.26** | **-15,410.43** | **0.00** | **0.10** |
| + | | + | + | + | | + | + | | | | 11 | 7,716.07 | -15,410.02 | 0.40 | 0.08 |
| + | + | + | | + | | + | + | | | | 11 | 7,715.74 | -15,409.37 | 1.05 | 0.06 |
| + | | + | | + | + | + | + | | | | 11 | 7,715.42 | -15,408.72 | 1.71 | 0.04 |
| + | | + | | + | | + | + | | | + | 13 | 7,717.40 | -15,408.65 | 1.78 | 0.04 |
| + | + | + | + | + | | + | + | | | | 12 | 7,716.34 | -15,408.54 | 1.89 | 0.04 |
| + | | + | + | + | | + | + | | + | | 12 | 7,716.33 | -15,408.52 | 1.90 | 0.04 |
| + | | + | + | + | | + | + | | | + | 14 | 7,718.33 | -15,408.46 | 1.96 | 0.04 |

Table S 6 Descriptive statistics of junco roost behavior by site

| | Hudson | Farhill | Dowler | Baker |
|---|---|---|---|---|
| **Variable** | **Mean ± SE (n)** | **Mean ± SE (n)** | **Mean ± SE (n)** | **Mean ± SE (n)** |
| Roost initiation (ET) | 17:31 ± 6.1min (98) | 17:45 ± 6.6min (93) | 17:22 ± 5.9min (97) | 17:57 ± 5.2min (105) |
| Roost initiation from sunset (hrs) | -1.12 ± 0.02 (98) | -0.82 ± 0.03 (93) | -1.15 ± 0.02 (97) | -0.67 ± 0.04 (105) |
| Departure time (ET) | 07:24 ± 1.8min (99) | 07:19 ± 2.5min (95) | 07:15 ± 2.1min (95) | 07:07 ± 2min (105) |
| Roost departure from sunrise (hrs) | 0.03 ± 0.02 (99) | -0.07 ± 0.03 (95) | -0.1 ± 0.02 (95) | -0.26 ± 0.02 (105) |
| Proportion time restless* | 0.01 ± 0 (96) | 0.01 ± 0 (88) | 0.01 ± 0 (88) | 0.02 ± 0 (104) |
| Restlessness (events/hr)* | 0.83 ± 0.04 (96) | 1.62 ± 0.16 (88) | 1.13 ± 0.14 (88) | 2.79 ± 0.19 (104) |
| Time roosting (hr)* | 13.79 ± 0.11 (96) | 13.37 ± 0.14 (88) | 13.97 ± 0.11 (88) | 13.01 ± 0.1 (104) |

Values are mean ± SE with sample size (number of nights observed) in parentheses.
* Summarized on smaller subset of data used for analyses of restlessness data where birds were observed > 40% of the night.

Table S 7 Descriptive statistics of environmental variables used in models of junco roost behavior by site.

| | Hudson | Farhill | Dowler | Baker |
|---|---|---|---|---|
| **Variable** | **Mean ± SE (n)** | **Mean ± SE (n)** | **Mean ± SE (n)** | **Mean ± SE (n)** |
| Evening temperature (°C) | 3.78 ± 0.85 (102) | 3.56 ± 0.87 (96) | 3.4 ± 0.91 (99) | 3.72 ± 0.84 (107) |
| Night temperature (°C) | 0.05 ± 0.9 (102) | -0.15 ± 0.92 (96) | -0.34 ± 0.94 (99) | 0.08 ± 0.89 (107) |
| Morning temperature (°C) | -0.51 ± 0.89 (102) | -0.73 ± 0.9 (96) | -0.72 ± 0.93 (99) | -0.41 ± 0.87 (107) |
| Night cloud cover (%) | 42.45 ± 3.56 (102) | 42.83 ± 3.67 (96) | 44.18 ± 3.68 (99) | 43.2 ± 3.47 (107) |
| Mean cloud cover (%) | 40.08 ± 2.74 (102) | 39.98 ± 2.81 (96) | 40.84 ± 2.84 (99) | 40.46 ± 2.66 (107) |
| Hours moon above horizon | 6.9 ± 0.37 (102) | 7.04 ± 0.4 (96) | 7.02 ± 0.38 (99) | 6.94 ± 0.36 (107) |
| Illumination (lux) | 14.8 ± 2.29 (102) | 15.76 ± 2.39 (96) | 13.9 ± 2.27 (99) | 13.11 ± 2.11 (107) |
| Log illumination | 0.97 ± 0.23 (102) | 1.33 ± 0.2 (96) | 0.62 ± 0.26 (99) | 0.76 ± 0.23 (107) |
| Afternoon precipitation (mm/hr) | 0 ± 0 (102) | 0 ± 0 (96) | 0 ± 0 (99) | 0 ± 0 (107) |
| Morning pressure (hPa) | 977.46 ± 0.82 (102) | 977.35 ± 0.85 (96) | 977.91 ± 0.81 (99) | 977.3 ± 0.79 (107) |
| Night pressure (hPa) | 976.77 ± 0.79 (102) | 976.7 ± 0.83 (96) | 977.34 ± 0.78 (99) | 976.72 ± 0.76 (107) |
| Mean pressure (hPa) | 977.01 ± 0.72 (102) | 976.9 ± 0.76 (96) | 977.52 ± 0.71 (99) | 976.92 ± 0.7 (107) |
| Morning wind speed (m/s) | 5.81 ± 0.24 (102) | 5.75 ± 0.25 (96) | 5.88 ± 0.25 (99) | 5.88 ± 0.24 (107) |
| Night wind speed (m/s) | 6.54 ± 0.32 (102) | 6.5 ± 0.33 (96) | 6.37 ± 0.31 (99) | 6.59 ± 0.31 (107) |
| Max wind speed (m/s) | 8.36 ± 0.28 (102) | 8.31 ± 0.28 (96) | 8.26 ± 0.28 (99) | 8.44 ± 0.27 (107) |
| Night cloud cover (%)* | 42.77 ± 3.7 (96) | 42.49 ± 3.82 (88) | 44.4 ± 3.92 (88) | 42.76 ± 3.56 (104) |
| Night precipitation (mm/hr)* | 0 ± 0 (96) | 0 ± 0 (88) | 0 ± 0 (88) | 0 ± 0 (104) |
| Night pressure (hPa)* | 976.76 ± 0.82 (96) | 976.52 ± 0.88 (88) | 977.17 ± 0.84 (88) | 976.78 ± 0.78 (104) |
| Night wind speed (m/s)* | 6.52 ± 0.33 (96) | 6.7 ± 0.34 (88) | 6.44 ± 0.33 (88) | 6.49 ± 0.31 (104) |

Values are mean ± SE with sample size in parentheses. ET = Eastern Time. * summarized on smaller subset of data used for analyses of restlessness data where birds were observed > 40% of the night.

Post hoc comparisions from emmeans.

Table S 2 Roost initiation

**Table R4. Pairwise comparisons of illumination slopes — Roost hour model**

| Contrast | Estimate | SE | df | z | p |
|---|---|---|---|---|---|
| Baker - Dowler | -0.0650 | 0.0095 | Inf | -6.8414 | **0.0000** |
| Baker - Farhill | -0.0570 | 0.0159 | Inf | -3.5879 | **0.0019** |
| Baker - Hudson | -0.0262 | 0.0097 | Inf | -2.6967 | **0.0353** |
| Dowler - Farhill | 0.0080 | 0.0160 | Inf | 0.5005 | 0.9590 |
| Dowler - Hudson | 0.0388 | 0.0099 | Inf | 3.9268 | **0.0005** |
| Farhill - Hudson | 0.0308 | 0.0161 | Inf | 1.9093 | 0.2240 |
